# NSD2 E1099K drives relapse in pediatric acute lymphoblastic leukemia by disrupting 3D chromatin organization

**DOI:** 10.1101/2022.02.24.481835

**Authors:** Sonali Narang, Nikki A. Evensen, Jason Saliba, Joanna Pierro, Mignon L. Loh, Patrick A. Brown, Heather Mulder, Ying Shao, John Easton, Xiaotu Ma, Aristotelis Tsirigos, William L. Carroll

## Abstract

The NSD2 p.E1099K (EK) mutation has been shown to be enriched in patients with relapsed ALL and found to play a role in clonal fitness dependent on the underlying genetic/epigenetic landscape of the cells. To uncover 3D chromatin architecture-related mechanisms underlying drug resistance, we systematically integrated Hi-C, ATAC-seq, RNA-seq and ChIP-seq data from three B-ALL cell lines heterozygous for NSD2 EK (RS4;11, RCH-ACV, SEM) and assessed changes upon knockdown. NSD2 knockdown revealed widespread remodeling of the 3D genome, specifically in terms of compartmentalization. Systematic integration of these datasets revealed significant switches in A/B compartments with a strong bias towards B compartments upon knockdown, suggesting that NSD2 EK plays a prominent role in maintaining A compartments through enrichment of H3K36me2 epigenetic marks. In contrast, we identified few changes in intra-TAD activity suggesting that the NSD2 EK impacts transcriptional changes through a remarkable dependence on compartmentalization. Furthermore, EK-mediated reorganization of compartments highlights the existence of a common core of compacting loci shared across the three cell lines that explain previously described phenotypes as well as serve as targets for therapeutic intervention. This study offers a novel mechanism by which NSD2 EK drives clonal evolution and drug resistance.

## INTRODUCTION

While the overall outcome with children with acute lymphoblastic leukemia (All) has improved dramatically, up to 20% of patients relapse making ALL one of the leading causes of cancer-related death in children ^1,2^. Genome-wide profiling studies have shown enrichment of mutations at relapse including somatic alterations in epigenetic regulators such as *MSH6*, *SETD2*, and *NSD2* ^3^. *NSD2* is a key histone methyltransferase involved in monomethylation and dimethylation of lysine 36 of histone 3 (H3K36), a mark associated with active transcription ^4^. Specifically, the recurrent gain of function mutation p.E1099K (EK) has been found to be enriched in patients with relapsed ALL ^5^.

The NSD2 EK mutation results in increased methyltransferase activity leading to a global increase in H3K36me2 levels (active mark) as well as the concomitant inhibition of EZH2-mediated H3K27me3 levels (repressive mark) in pediatric ALL ^6,7^. Our previous studies have shown knockdown of NSD2 in EK mutated B-ALL cell lines resulted in decreased proliferation, decreased clonal growth, and increased sensitivity to cytotoxic chemotherapeutic agents with no effect on NSD2 wildtype lines ^5^. Other work also demonstrated changes in growth and clonogenicity as well as cellular adhesion upon CRISPR mediated reversion of EK to wildtype in NSD2 mutated lines ^7^. Interestingly, RNA-seq data from both studies revealed variable transcriptional reprogramming upon loss of NSD2 in different cell line models. While some common pathways were identified, such as cell adhesion and Rap1 signaling, collectively the data suggests cell context specific changes occur in response to mutated NSD2. Importantly, we also demonstrated minimal overlap in chromatin accessibility changes upon NSD2 knockdown in the three EK harboring cell lines ^5^. Our current work addresses the transcriptional and chromatin accessibility heterogeneity observed as a result of the NSD2 EK mutation.

In this study, we investigate the role 3D genome organization plays in EK-mediated relapse. 3D genome organization refers to the strategic positioning of regulatory elements to regions best suited for the regulation of genome function. The organizational hierarchy is made up of multiscale structural units such as chromosomal territories, A/B compartments, topologically associating domains (TADs), and chromatin loops each of which play an important role in regulating gene expression ^8,9^. At the Mb scale, chromosomes are spatially divided into two major domains, A and B compartments, that correspond to active and inactive chromatin, respectively ^10,11^. At the sub-Mb scale, the genome can be further subdivided into highly self-interacting chromatin units referred to as TADs ^10,12,13^. TADs play a major role in the regulation of gene expression by restricting the influence of regulatory elements to genes within the same TAD as well as insulating them from interactions with neighboring domains ^14^.

Dysregulation of higher-order genomic architecture in several disease models has been linked to changes in the epigenetic landscape ^15–18^. In multiple myeloma, expansion of H3K26me2 and shrinkage of H3K27me3 domains as a result of NSD2 overexpression were shown to be linked to chromatin changes in TADs and CTCF loops as well as disruptions in gene expression ^15^. In lymphoma, profound decompaction of the genome as a result of disruption in H1 function was shown to drive changes in the epigenetic landscape including the gain of H3K36me2 and loss of H3K27me3 leading to the aberrant expression of normally silenced stem cell associated genes ^16^. In B-ALL, however, NSD2 mediated context dependent phenotypic changes have yet to be linked to the dysregulation of higher-order structures.

To uncover 3D chromatin architecture-related mechanisms underlying NSD2 mediated drug resistance and disease progression, here we performed Hi-C and systematically integrated it with previously published ATAC-seq, RNA-seq, and ChIP-seq data from three B-ALL cell lines heterozygous for NSD2 EK (RS4;11, RCH-ACV, SEM) and assessed changes upon knockdown. Systematic integration of these datasets revealed significant switches in A/B compartmentalization with a strong bias towards B compartments suggesting that NSD2 EK plays a prominent role in maintaining A compartments through enrichment of the H3K36me2 epigenetic mark. We also identified few changes in intra-TAD activity suggesting that the NSD2 EK exemplifies a remarkable dependence on compartmentalization. Furthermore, EK-mediated reorganization of compartments highlights the existence of a common core of compacting loci shared across the three cell lines that could explain previously described shared phenotypes such as proliferation, as well as serve as targets for therapeutic intervention. This study offers a novel mechanism by which NSD2 EK drives ALL clonal evolution in a cell context dependent manner.

## RESULTS

### NSD2 knockdown drives widespread changes in A/B compartments

We performed Hi-C and systematically integrated it with previously published ATAC-seq, RNA-seq, and ChIP-seq data from three B-ALL cell lines heterozygous for NSD2 EK (RS4;11, RCH-ACV, SEM) either expressing a non-targeting shRNA or an NSD2 targeting shRNA henceforth referred to as NSD2 High and Low cells ^5^ (Fig 1a). Hi-C was performed using the Arima Kit and Hi-C data was processed by our HiC-bench platform and showed alignment rates with a high number of usable intra-chromosomal long-range read pairs (~100 million) ^19^ (Supplementary Fig. 1a). We first assessed compartmentalization with the CscoreTool algorithm by calling A/B compartments ^20^. Principal Component Analysis (PCA) of Cscore compartment scores revealed a separation between NSD2 High and Low cell lines (Fig. 1b). Cscore compartment calls showed similar percentages of A/B compartments in NSD2 High and NSD2 Low cell lines (Fig. 1c). We next examined changes in A/B compartments upon NSD2 knockdown. We observed ~7.49% compartment switching between NSD2 High and NSD2 Low across the three cell lines (Fig. 1d). Overall, a greater number of compartment switches occurred from A to B upon knockdown with 3,047 regions switching from A to B and 1,132 regions switching from B to A. Of the switching compartments, we identified 226 shared switches across the three cell lines from A to B and 64 from B to A (Fig. 1e). 30.39% (926/3047) of A to B switches and 42.93% (486/1132) B to A switches were shared by at least 2 cell lines (Fig. 1f). One such compartment switch from A to B shared by all three cell lines involved the Neogenin 1 (NEO1) gene locus, which has been implicated in cell adhesion (Fig. 1g left panel) ^21^. Additionally, we identified cell-line specific switches including an A to B switch at the PR Domain Zinc Finger Protein 8 (PRDM8) and FGF5 gene locus in RS4;11 (Fig. 1g right panel).

**Figure 1.**
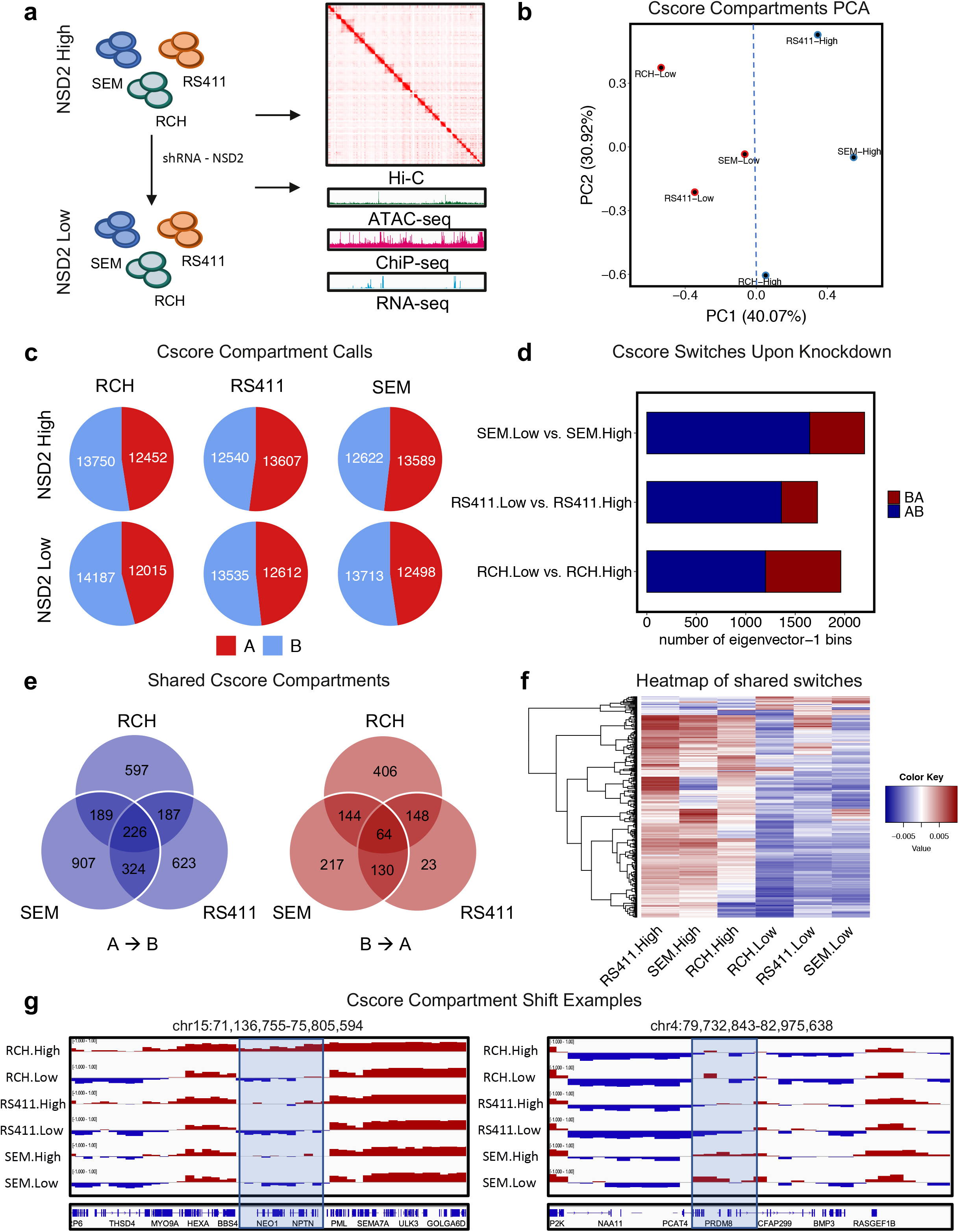
NSD2 EK knockdown drives A/B compartment reorganization. **a.** Schematic demonstrating overall study design. **b.** PCA of A/B compartment calls with CScore. **c.** Pie chart showing numbers of A and B compartment calls for each cell line in NSD2 High and Low cell lines. **d.** Bar plot showing number of compartment switches for each cell line upon knockdown. **e.** Venn diagram showing overlap of switching compartments between three B-ALL cell lines upon knockdown, A to B and B to A switches (left and right respectively). **f.** Heatmap representation of compartment shifts shared by at least 2 cell lines upon knockdown. **g.** IGV tracks of an example of A to B compartment shift shared by all three cell lines at the NEO1 locus (left). IGV tracks of an example of a compartment shift specific to RS4;11 cell line at the PRDM and FGF5 locus (right).

### NSD2 knockdown-related compartment switches alter gene expression in a cell-context dependent manner

To understand how NSD2 knockdown alters gene expression, we first analyzed previously published RNA-seq data from NSD2 High and NSD2 Low cell lines ^5^. PCA revealed that NSD2 High and NSD2 Low replicates separated into distinct cell-type specific clusters demonstrating transcriptional heterogeneity (Fig. 2a). As previously described, we observed that NSD2 knockdown leads to the deregulation of several genes across all three cell lines (Fig. 2b). NSD2 was notably confirmed to be downregulated upon NSD2 knockdown amongst other genes (Fig. 2b). Overlap analysis also revealed changes in gene expression to be predominantly cell-context dependent with minimal overlap between cell lines (Supplementary 2a). Only 1.24% of differentially expressed genes were shared between the three cell lines.

**Figure 2.**
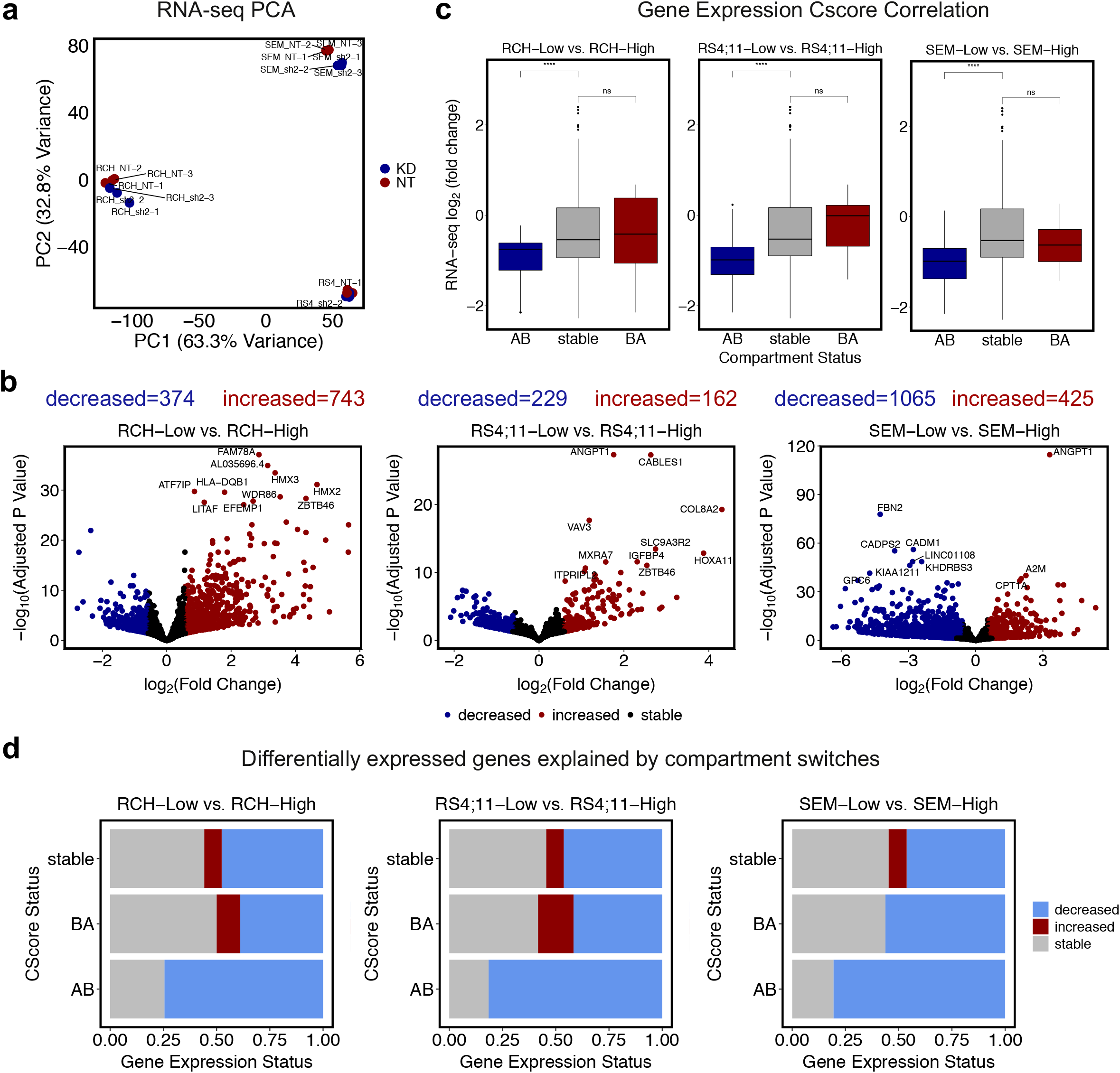
NSD2 knockdown-related A to B compartment switches correlate with gene expression changes. **a.** PCA representation of triplicate RNA-seq data showing top 1000 genes. **b.** Correlation boxplots of gene expression changes and compartment switch status upon knockdown (AB, BA, stable) for each cell line. **c.** Volcano plots demonstrating differentially expressed genes (abs(L2FC) > 0.58, p-value < 0.05) upon knockdown per cell line with top 10 labeled. **d.** Association barplots showing fraction of genes (increased, decreased, stable) at compartment shifts (AB, BA, stable).

To examine the relationship between alterations in A/B compartments and gene expression, compartment switches were categorized as A to B, B to A, or stable and then assessed for changes in gene expression. AB compartment switches significantly correlated with downregulated genes; however, BA compartment switches did not correlate with upregulated genes (Fig. 2c). Likewise, we observed that the A to B compartment switches were characterized by a significant fraction of downregulated genes whereas B to A compartment switches did not show such a relationship with upregulated genes (Fig. 2d). This data suggests that EK knockdown is responsible for gene expression changes specifically through compartment reorganization from A to B.

To expand our analysis, we incorporated compartment shifts in addition to compartment switches (Fig. 3a). Shifts include A to more A, B to more B, A to less A, and B to more B. The addition of compartment shifts revealed 26.50% compartment changes between NSD2 High and NSD2 Low cell lines (Fig. 3b). Further exploring concordance with gene expression revealed that compartment changes significantly correlated with gene expression changes (Fig. 3c). We also identified that the majority of A to B switches and shifts were made up of significantly downregulated genes (Fig. 3d; L2FC < 0.58, p-value < 0.05), whereas B to A switches and shifts did not demonstrate a clear pattern similar to our findings with switches alone (Fig. 3d, Supplementary Fig. 2c).

**Figure 3.**
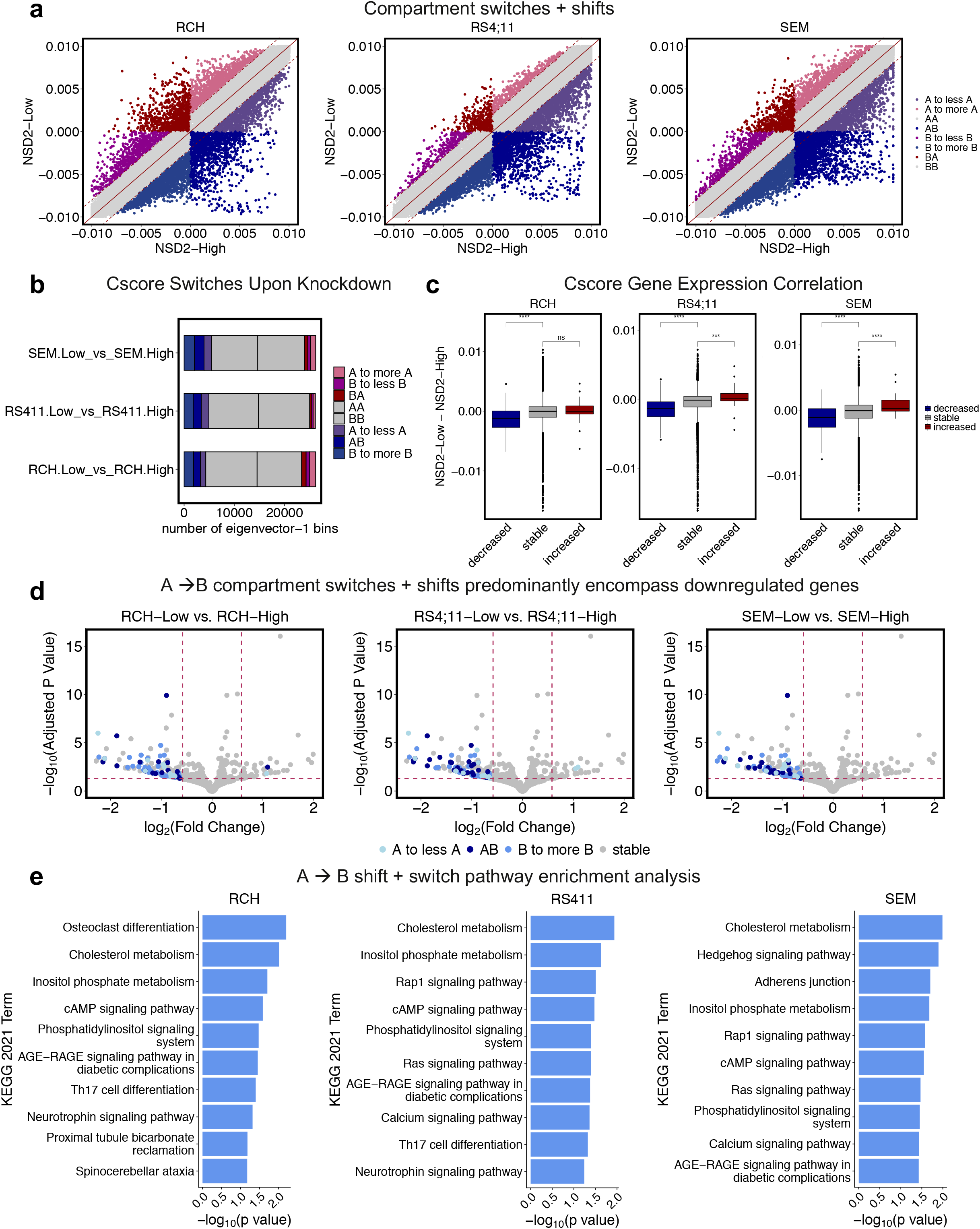
NSD2 knockdown-related A to B compartment switches and shifts correlate with gene expression changes. **a.** Scatter plot demonstrating strategy for calling 6 categories of compartment switches and shifts (A to B, and B to A compartment switches; A to less A, A to more A, B to less B, and B to more B compartment shifts). **b.** Bar plot showing number of compartment switches and shifts for each cell line upon knockdown. **c.** Correlation boxplots of compartment cscore and gene expression changes upon knockdown (abs(L2FC) > 0.58, p-value 0.05) for each cell line. **d.** Volcano plots demonstrating differentially expressed genes (abs(L2FC) > 0.58, p-value < 0.05) highlighted by compartment switch or shift (A to less A, A to B, B to more B, or stable). **e.** KEGG 2021 pathway enrichment analysis of genes differentially expressed (L2FC < −0.58, p-value < 0.05) at A to B compartment switches or A to less A and B to more B compartment shifts per cell line.

To further explore this concordance between compartment and gene expression changes, we assessed the number of differentially expressed genes explained by compartment changes. Approximately 7.60% of genes downregulated upon knockdown could be explained by compartments that switched from A to B (Supplementary Table 1). With the addition of compartment shifts, 20.00% of downregulated genes could be explained by compartment switches and shifts (Supplementary Table 1). In contrast, only 1.20% of upregulated genes could be explained by compartment changes. Ascribing a concordance score to each compartment shift and switch revealed that concordance was observed specifically in those compartments that compacted (Supplementary Fig. 2d).

Lastly, to better understand the impact these changes have on downstream signaling, we performed pathway enrichment analysis for those differentially expressed genes that were associated with either a compartment switch or shift per cell line (Fig. 3e). Interestingly, cancer-related pathways previously identified by RNA-seq or ChIP-seq analysis ^5,22^ were also found in our compartment-based analysis including Rap1, Ras, Phosphatidylinositol 3-kinase, and calcium signaling. Furthermore, these pathways were shared by at least two of the three cell lines providing evidence that the existence of a core of compacting loci can explain previously described shared phenotypes such as proliferation ^5^.

### NSD2 knockdown-related compartment switches associate with chromatin accessibility changes

To explore how NSD2 knockdown alters chromatin accessibility, ATAC-seq was performed on the NSD2 High and NSD2 Low cell lines. As previously described ^5^, NSD2 knockdown leads to the restructuring of a modest number of peaks across all three cell lines. This data showed a significant loss of ATAC-seq peaks for two of the three cell lines whereas SEM showed a paradoxical bias towards increased accessibility (Fig. 4a). PCA and heatmap revealed that NSD2 High and NSD2 Low replicates separated into distinct cell-type specific clusters demonstrating chromatin accessibility heterogeneity (Supplementary Fig. 3a,b). Overlap analysis also revealed changes to be predominantly cell-context dependent with minimal overlap between cell lines (Supplementary 3c). Additionally, we observed a shift in the distribution of the ATAC-seq peaks from intergenic regions towards promoter regions as has been previously shown with H3K36me2 peak distribution (Fig. 4b) ^7,23^.

**Figure 4.**
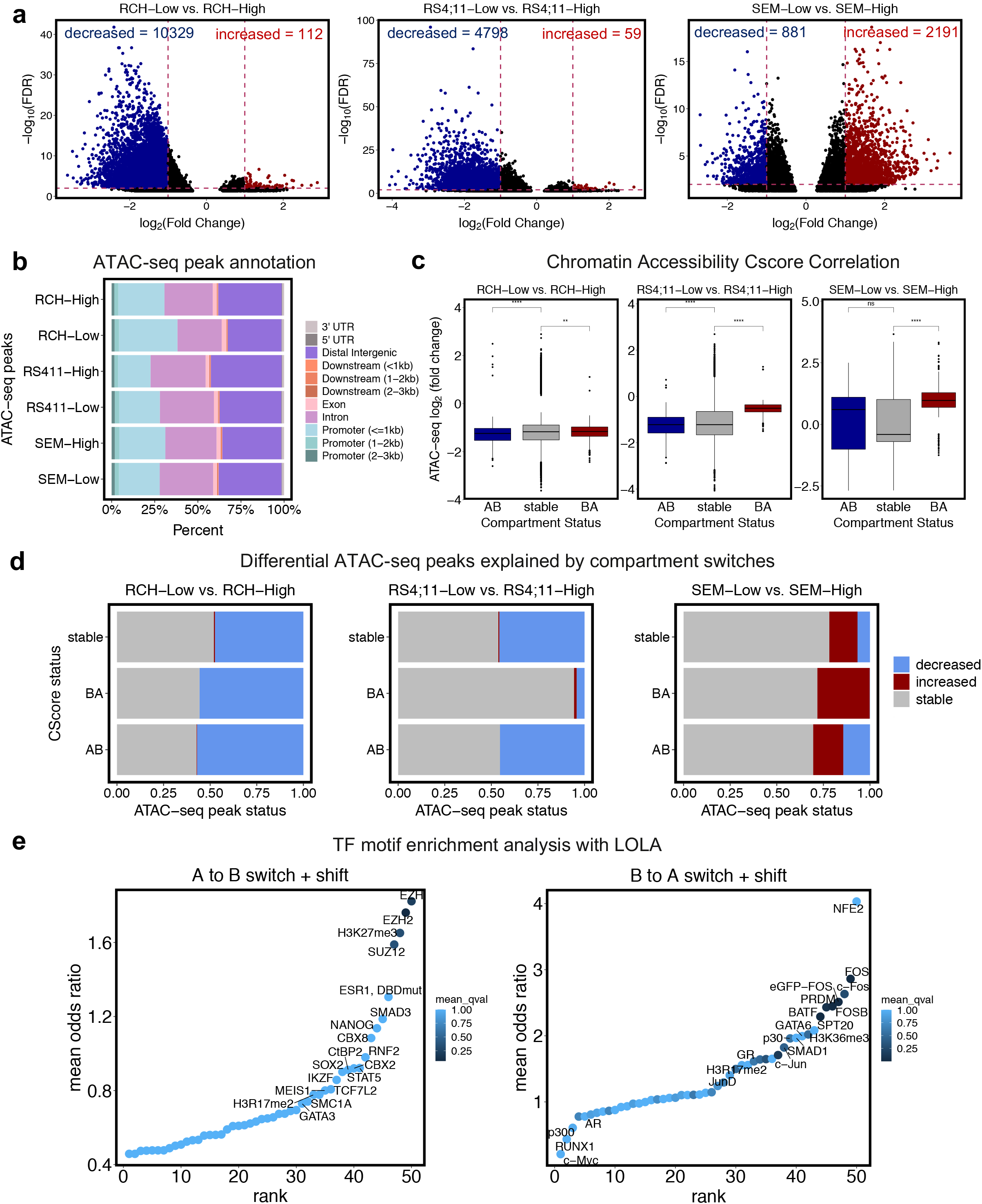
NSD2 knockdown leads to cell-type specific alterations in chromatin accessibility. **a.** Volcano plots demonstrating differentially accessible regions upon knockdown per cell line (abs(L2FC) > 1, FDR < .01). **b.** Bar plot showing distribution of annotated ATAC-seq peaks for each NSD2-High and NSD2-Low cell line. **c.** Correlation boxplots of chromatin accessibility changes and compartment switches (AB, BA, stable) for each cell line. **d.** Association barplots showing fraction of ATAC-seq peaks (increased, decreased, stable) at compartment switches (AB, BA, stable). **e.** TF motif enrichment analysis using LOLA for ATAC-seq peaks concordant with A to B switches and shifts (left) and B to A switches and shifts (right).

To examine the relationship between alterations in A/B compartments and chromatin accessibility, compartment switches were categorized as A to B, B to A, or stable and then assessed for changes in accessibility. Chromatin accessibility data did not show correlation with compartment data across the three cell lines. The majority of differential peaks were decreased regardless of directionality of compartment changes for RCH-ACV and RS4;11 lines. The opposite was true for SEM, which showed increased accessibility overall (Fig. 4c). Compartment switches were characterized by predominantly stable ATAC-seq peaks. We also noticed significant variability in the association with differential peaks across the three cell lines (Fig. 4d).

Exploring this link further, we identified concordant changes in A/B compartments, gene expression, and chromatin accessibility at the NEO1 and PRDM8/FGF5 loci. Along with the shared compartment switch from A to B in all three cell lines at the NEO1 locus, we also identified concordant decreases in chromatin accessibility and gene expression shown by genome browser tracks (Supplementary Fig. 3e). Similarly, along with the RS4;11 cell-type specific compartment switch from A to B at the PRDM8/FGF5 locus, we identified concordant decreases in chromatin accessibility and gene expression in RS4;11 cells shown by genome browser tracks (Supplementary Fig. 3f).

To identify key regulators of transcription attributed to compartment switches, we performed a transcription factor (TF) motif enrichment analysis with concordant increased or decreased ATAC-seq peaks, gene expression, and compartment switches and shifts (Fig. 4e). EZH2 and H3K27me3, involved in repression, were among the top hits for sites with A to B compartment changes. Notably, factors linked to stem cell functionality, including NANOG, SUZ12, SOX2, and Meis1 were also enriched ^24^.

### NSD2 knockdown results in few intra-TAD activity changes

Following our compartment analysis, we investigated TAD activity changes between NSD2 High and NSD2 Low cell lines. Assessing mean TAD activity across samples revealed that the bulk of TADs remain stable upon knockdown across the three cell lines (Fig. 5a). Comparison of intra-TAD activity between NSD2 High and NSD2 Low cell lines identified several statistically significant decreases and very few increases across all three cell types (Fig. 5b, Supplementary Fig. 4a; FDR < 0.1 and abs(L2FC) > 0.25). Of the differential TADs, we identified no shared increases in TAD activity and 7 shared decreases in TAD activity between the three cell lines suggesting that changes in TAD activity were cell-context dependent (Fig. 5c).

**Figure 5.**
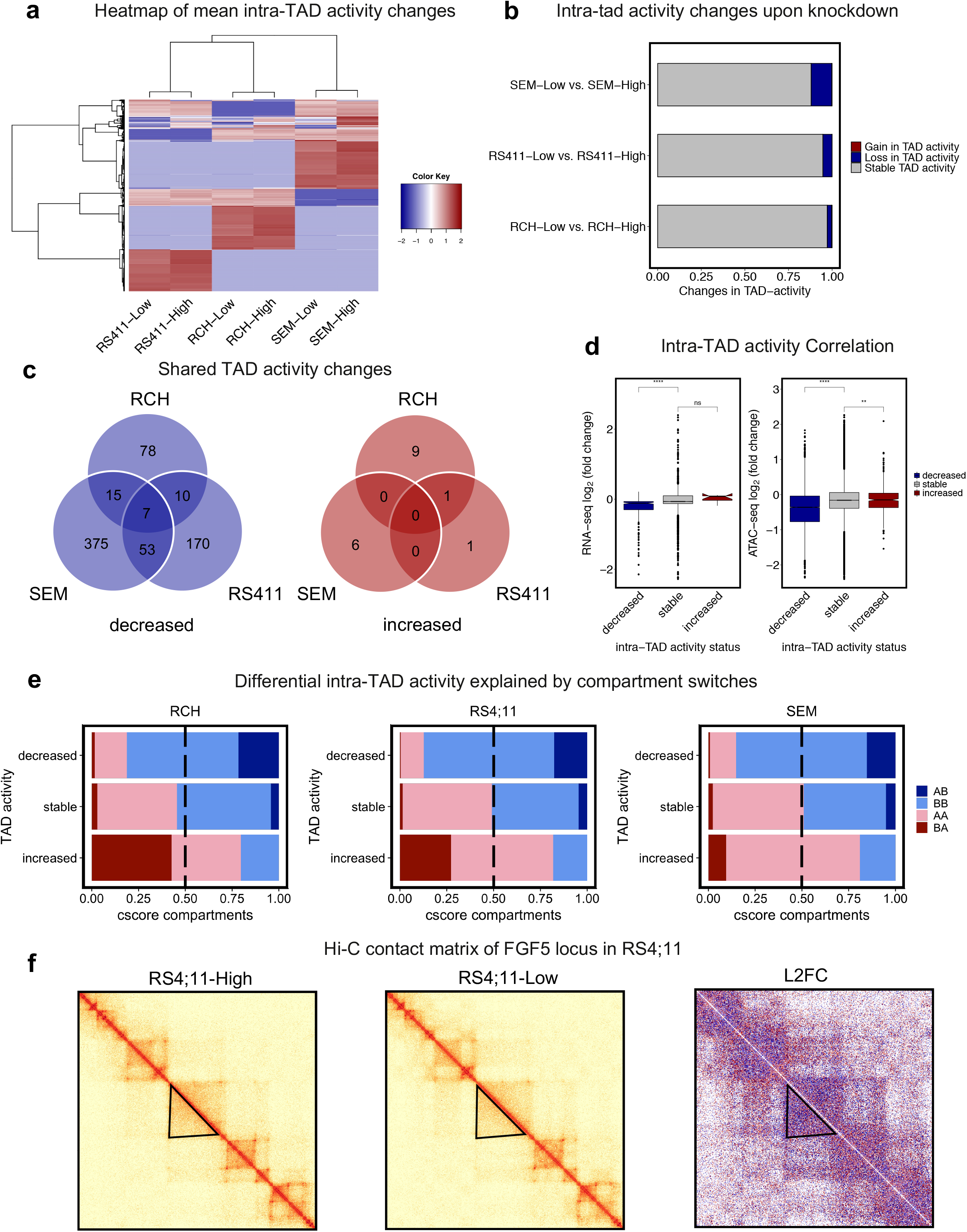
NSD2 knockdown has minimal impact on intra-TAD activity. **a.** Heat map representation of mean intra-TAD activity per cell line. **b.** Bar plot showing number of TAD activity changes per cell line upon knockdown (abs(L2FC) > 0.25, FDR < 0.01). **c.** Venn diagram showing overlap of differential TADs between three B-ALL cell lines, decreased and increased (left and right respectively). **d.** Correlation boxplots showing gene expression and chromatin accessibility changes within differential TADs upon NSD2 knockdown. **e.** Association barplots showing fraction of intra-TAD activity changes(increased, decreased, stable) at compartment switches (AB, BA, AA, BB). **f.** Example of a TAD activity alteration using Hi-C contact matrices for RS4;11 at the FGF5 locus showing a decrease in intra-TAD activity (NSD2-High, NSD2-Low, and L2FC(Low/High) from left to right).

To investigate whether changes in intra-TAD activity associated with changes in gene expression and chromatin accessibility, we performed differential expression analysis (abs(L2FC) >0.58, p <0.05) and differential chromatin accessibility analysis (abs(L2FC) >0.32, FDR <0.01) within differentially active TADs. Although few differential TADs were identified, integration of gene expression changes and chromatin accessibility changes with differentially active TADs indicated a significant correlation in both cases (Fig. 5d). In addition, we observed that increases in TAD activity predominantly occurred in regions with B to A compartment switches and decreases in TAD activity predominantly occurred in regions with A to B compartment switches (Fig. 5e). In addition to earlier data, we identified a cell-type specific decrease in TAD activity at the FGF5 locus in RS4;11 concordant with compartment switch, chromatin accessibility, and gene expression as shown by the contact matrix at NSD2 High, NSD2 Low, and L2FC contact matrix (Low/High) (Fig. 5f).

### NSD2 mutant patient samples reflect B-ALL cell line data

To examine how relapse enriched NSD2 EK mutation behaves in patient samples, we acquired expression data from three matched diagnosis (NSD2 Low) and relapse (NSD2 High) patient pairs (Supplementary Table 2). PCA revealed that NSD2 High samples separated from the NSD2 Low samples by the first principal component (Fig. 6a). Heatmap and volcano plots both demonstrate significant NSD2-mediated changes in gene expression with 525 decreased and 514 increased genes (Fig. 6b,c; abs(L2FC) >0.58, p-adj <0.05). To test our hypothesis that EK-mediated reorganization of compartments affects gene expression in relapse, we compared gene expression data from the patient samples to expression and compartment data from the three cell lines. Comparing differentially expressed genes that were downregulated in NSD2 Low cell lines to those downregulated in NSD2 Low patients revealed significant overlap whereas, as expected, cell line NSD2 Low downregulated genes with patient NSD2 Low upregulated genes did not (Supplementary Fig. 5a). Comparing those cell line NSD2 Low genes downregulated and mediated by compartment switches and shifts to those downregulated in NSD2 Low patients also revealed significant overlap (Fig. 6d). Additionally, differentially expressed genes in patients were found to be enriched in A to B changes compared with B to A changes (Fig. 6e). Downregulated genes in NSD2 Low patients were also found to correlate with A to B changes whereas upregulated genes did not correlate with B to A changes as we had observed in cell lines (Fig. 6f). Lastly, we performed pathway enrichment analysis with those genes downregulated in NSD2 Low patients that overlapped cell line compartment changes (Fig. 6g). Using this patient data, we identified numerous pathways that have been shown previously with NSD2 cell line models, such as calcium signaling, MAPK, and cell adhesion pathways.

**Figure 6.**
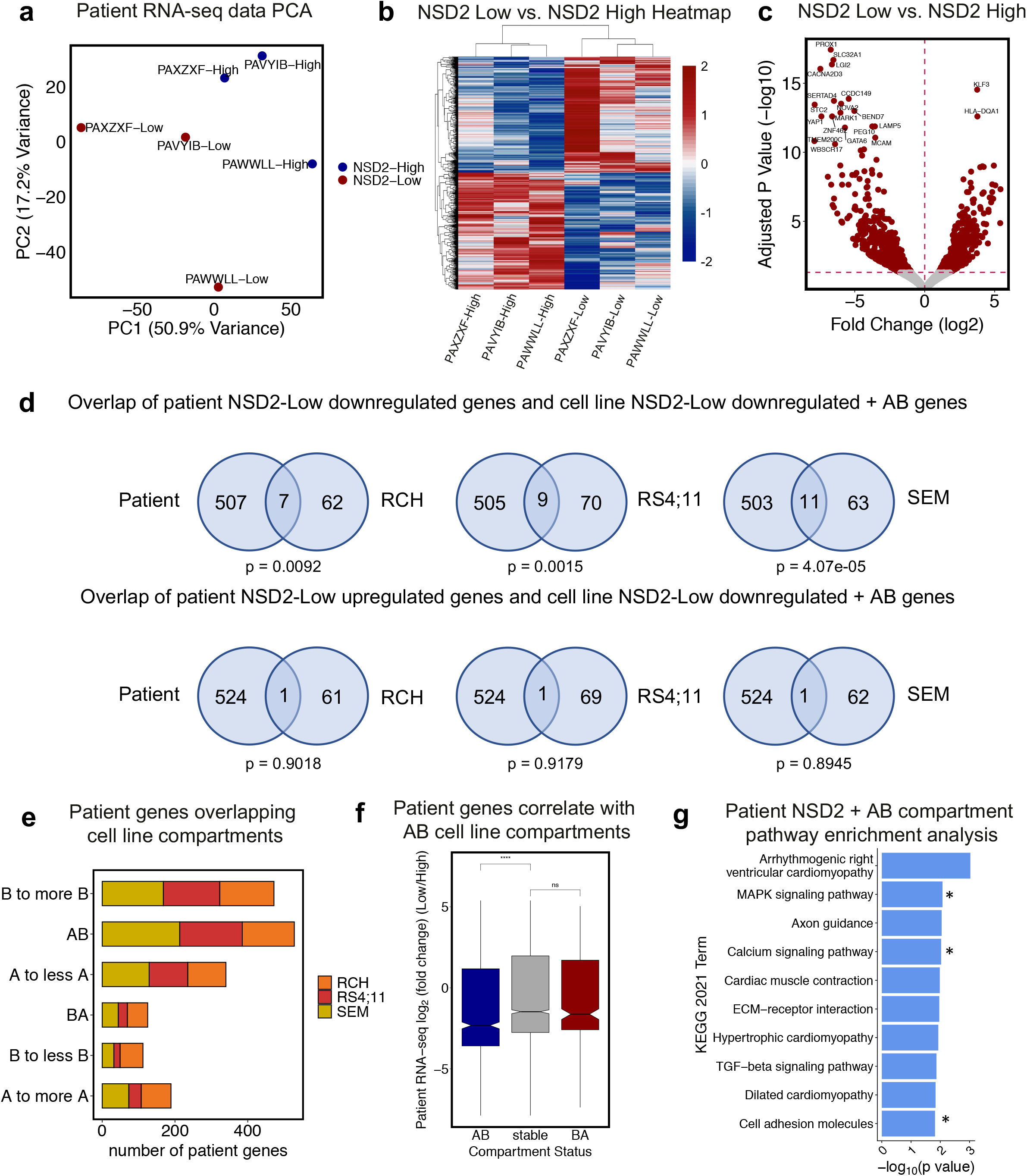
NSD2 mutant patient samples reflect B-ALL cell line data. **a.** PCA of 3 matched diagnosis (NSD2-Low) and relapse (NSD2-High) patient pair gene expression data. **b.** Heatmap of significantly expressed genes upon relapse (abs(L2FC) > 0.58, p-adjusted < 0.05). **c.** Volcano plot demonstrating differentially expressed genes highlighted in red(abs(L2FC) > 0.58, p-adjusted < 0.05). **d.** Venn diagrams demonstrating significant overlap of patient NSD2-Low downregulated genes and cell line NSD2-Low downregulated genes with AB compartment switches and insignificant overlap of patient NSD2-Low upregulated genes and cell line NSD2-Low downregulated genes with AB compartment switches (one-tailed Fisher’s exact test). **e.** Barplot demonstrating number of patient genes overlapping cell line compartment switches and shifts colored by cell line. **f.** Correlation boxplot showing patient gene expression changes within cell line compartment changes (NSD2-Low/NSD2-High). **g.** KEGG 2021 pathway enrichment analysis of patient genes downregulated in NSD2-Low patients (abs(L2FC) > 0.58, p-adjusted < 0.05) that overlap with cell line A to B compartment switches, A to less A, and B to more B compartment shifts from any of the three cell lines. Starred pathways represent pathways previously shown in NSD2 models.

### NSD2 knockdown-mediated epigenetic changes correlate with 3D genome changes

We previously performed ChIP-seq on the RS4;11 NSD2 High and NSD2 Low cell lines with 1368 total significant changes for H3K36me2, and more modest changes for H3K27me3 and H3K27ac 5 (Fig. 7a). As noted previously H3K36me2 distribution shifted from predominantly intergenic regions to promoter regions upon NSD2 knockdown. Conversely, H3K27me3 peaks shift from promoter regions to intergenic regions upon knockdown (Fig. 7b). Notably, H3K27ac was found to be enriched at promoter regions with little difference in distribution upon knockdown.

**Figure 7.**
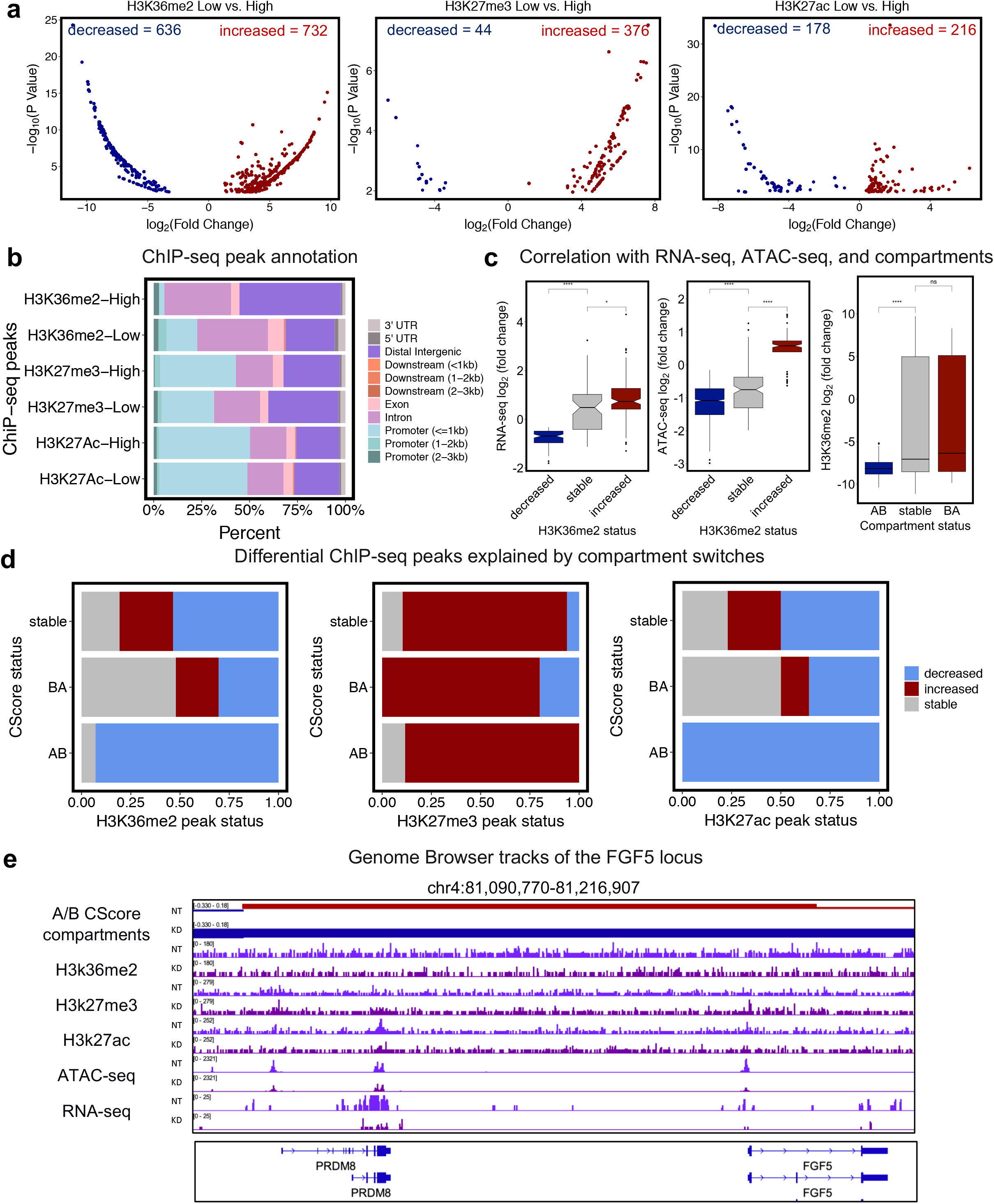
NSD2 knockdown-mediated epigenetic changes correlate with 3D genome changes. **a.** Volcano plots demonstrating significantly differential chip-seq peaks. **b.** Bar plot showing distribution of annotated ChiP-seq peaks for each epigenetic mark for NSD2-High and NSD2-Low RS4;11 cell line. **c.** Correlation boxplots of gene expression and chromatin accessibility changes with H3K36me2 chip-seq changes, and H3K36me2 alterations with cscore compartment switches. **d.** Association barplots showing fraction of differential chip-seq peaks (H3K36me2, H3K27me3, and H3K27ac) at cscore compartment switches (AB, BA, and stable). **e.** IGV genome browser tracks showing an example of concordant changes in compartments, chip-seq, gene expression, chromatin accessibility upon knockdown at the FGF5 gene locus.

To examine the relationship between alterations in the distribution of these marks and gene expression, chromatin accessibility, and compartments, we categorized the H3K36me2 peaks into those that are increased, decreased, or stable and assessed both gene expression and chromatin accessibility changes. H3K36me2 peaks were positively correlated with gene expression changes and chromatin accessibility (Fig. 7c). Compartment switches were also categorized as AB, BA, or stable and then assessed for changes in H3K36me2. H3K36me2 peaks positively correlated with AB switches but did not correlate with BA switches (Fig. 7c). We also identified that A to B compartment switches were associated with decreased H3K36me2 and H3K27ac peaks whereas B to A switches were associated with increased H3K27me3 peaks (Fig. 7d). Lastly, 14.30% of ChIP-seq peaks could be explained by A to B compartment changes (Supplementary Table 3). This data suggests that depletion of H3K36me2 as a result of NSD2 knockdown provides a more compact chromatin landscape.

Returning to the PRDM8/FGF5 locus, we saw decreased H3K36me2 peaks, increased H3K27me3 and H3K27ac peaks across the region along with an AB compartment switch (Fig. 7e). This example clearly shows the relationship between NSD2 EK-mediated epigenetic modifications and 3D genome organization.

## DISCUSSION

The NSD2 EK mutation is enriched in patients that relapse with ALL indicating a role in clonal evolution. Additionally, *in vitro* data confirms roles in cell proliferation, clonogenicity, and therapy resistance. We and others have demonstrated that the phenotypic impact of NSD2 EK is highly cell context dependent supporting a role in providing plasticity to overcome the selective pressures of therapy. Here we show that NSD2 EK plays an essential role in reorganizing the 3D genome, explaining the pleotropic effects observed to date, through a striking reliance on compartment reorganization.

Hi-C analysis of NSD2 High and NSD2 Low cell lines revealed significant compartment changes (26.50%). This is consistent with a multiple myeloma model in which 23.00% of compartments changes were identified ^15^. This is also in contrast to the 3.00-5.00% of changes identified in various stages of B-cell differentiation including pre-B to pro-B and pro-B to immature-B (unpublished data). Consistent with NSD2 EK’s role in increased H3K36me2 methyltransferase activity, we observed significantly more A to B compartment switches and shifts upon knockdown, compared to B to A shifts. This is in opposition to the multiple myeloma model in which low expressing vs high expressing cells showed comparable A to B and B to A switches^15^.

Importantly, those compartments that switched to a less compact state showed decreased H3K36me2 and increased H3K27me3 epigenetic marks, suggesting that NSD2 EK plays a prominent role in maintaining A compartments. The expansion of H3K36me2 past gene bodies into more distal intergenic regions was also lost upon knockdown of NSD2. These data suggest that NSD2 EK creates a more open, active chromatin environment by spreading the H3K36me2 mark perhaps endowing cells with an increased repertoire of genes to respond effectively to the selective pressures of treatment.

Furthermore, compartment changes and epigenetic changes significantly correlated with genes that were downregulated upon NSD2 knockdown. For those compartments that became more active (B to A) upon knockdown, we did not see a clear association with increased gene expression. Whether alterations in the epigenetic landscape can impact chromosome organization in a manner that corresponds to alterations in gene expression has been a long-standing question in the field. As Lhoumaud et al. described, these data together also highlight a bidirectional relationship between 2D and 3D chromatin structure in gene regulation ^15^.

Although EK-mediated changes in transcriptional output as well as chromatin accessibility differed dramatically between cell lines, pathway analysis on differential genes with compartments that changed from A to B demonstrated convergence on similar pathways, such as cell adhesion, Ras, and calcium related pathways. Interestingly, these pathways have also come up in our previous analysis with RNA-seq data alone as well as in other models ^7^. In addition, WNT and MAPK (Ras-related) pathways were identified in two out of the three lines examined and have also previously been linked to NSD2 in the multiple myeloma model ^15^. Our RNA-seq data from 3 patient pairs harboring the EK mutation at relapse also support this hypothesis as we saw significant overlap with the NSD2 EK cell lines. The identification of common downstream targets among preclinical models and patient samples has important clinical implications for possible future therapeutic interventions.

By examining the regions where we observed compartment compaction, decreased accessibility, and downregulation of gene expression upon knockdown of NSD2, we identified enrichment of TF motifs linked to stem cell functionality, including NANOG, SUZ12, SOX2, and Meis1. In the B-cell lymphoma model, Yusufova et al. also demonstrated that many differentially expressed genes were linked to stem cell functionality as well as enriched for various stem cell signature related pathways ^16^. This data suggests a potential core transcription factor network that may be dependent on compartmentalization.

In contrast to the identification of widespread compartment switches upon NSD2 knockdown, we identified very few changes in intra-TAD activity. This data suggests that NSD2 EK mediated changes are primarily linked to changes in compartmentalization. Furthermore, this supports the notion that TADs and compartments are established by separate, competing mechanisms. Our findings are also in agreement with a recent paper demonstrating H1 loss-of-function mutations in B-cell lymphomas leading to chromatin decompaction. Few TAD changes were observed and H1 loss led to decompaction through increased H3K36me2, again supporting the idea of NSD2 maintaining an open state ^16^. Although few TAD activity changes were identified, decreasing TAD activity was predominantly associated with A to B compartment switches and increasing TADs predominantly associated with B to A switches suggesting that TADs might be susceptible to patterns established by compartments.

Here we have identified a link between NSD2 EK mediated epigenetic changes and dysregulation of higher-order genomic architecture in B-ALL. Our study revealed NSD2 EK’s prominent role in maintaining A compartments through enrichment of H3K36me2 epigenetic marks. We also demonstrated NSD2 EK’s remarkable reliance on compartment reorganization over TAD activity. Despite transcriptional and chromatin accessibility heterogeneity across the three cell lines, this study highlights the existence of a common core of compacting loci that can explain previously described universal features, such as proliferation, as well as serve as targets for therapeutic intervention. Ultimately, this study offers a novel mechanism by which NSD2 EK drives ALL relapse by maintaining an open chromatin state to allow for transcriptional reprogramming in response to selective pressures associated with treatment.

## MATERIALS AND METHODS

### Experimental Procedures

Cell lines, RNA-seq, ATAC-seq, and ChIP-seq methods published previously ^5^.

### Primary patient sample experimental procedures

Cryopreserved PBMC samples from Children’s Oncology Group (COG) B-ALL study AALL-1331 were obtained from the COG Biobank. All subjects provided consent for banking and future research use of these specimens in accordance with the regulations of the institutional review boards of all participating institutions. Samples were resuspended in TRIzol reagent (Life Technologies) then processed using 5prime Phase Lock Gel Heavy tubes (Quantabio) following the manufacturer’s instructions. Following precipitation, the RNA was quantitated using the Qubit RNA BR assay kit (Life Technologies) and evaluated for quality using the Agilent 2100 Bioanalyzer with the RNA nano chip (Agilent). The KAPA RNA HyperPrep Kit with RiboErase (HMR) (Roche) with 100-200ng of total RNA was used for library construction with either Nextflex-96 RNA barcode adapters (BIOO Scientific, Illumina HiSeq 4000 platform), or IDT for Illumina unique dual indices adaptors (Illumina, Novaseq 6000 platform) and 12 PCR cycles of library amplification. Libraries were evaluated on the Bioanalyzer 2100 using the DNA 1000 chip. Libraries were sequenced using either the HiSeq 4000 (Illumina) or NovaSeq 6000 (Illumina) paired end 2X150 cycle protocol.

### Hi-C experimental procedures

Five million cells from actively growing NSD2 High and Low cell lines were crosslinked with 2% PFA and quenched with glycine. Following crosslinking, cells were lysed, DNA digested and proximally ligated following Arima Genomics’ Hi-C kit User Guide for Mammalian Cell Lines by the NYU Langone Genome Technology Center. Hi-C libraries were generated using Arima HiC Kit along with Swift 2S Plus Kit and Swift Indexing Kit following Arima-HiC Kit Library Preparation using Swift Biosciences(R) Accel-NGS(R) 2S Plus DNA Library Kit User Guide. Libraries were sequenced using Illumina’s NovaSeq 6000.

### Computational Analysis

#### ATAC-seq analysis

Cell line ATAC-seq data was processed with two replicates. Paired-end reads were aligned to the human reference genome(hg19) with Bowtie2 (v2.3.4.1) ^25^. Reads with a mapping quality <30 were removed. Duplicated reads were removed using Sambamba (v0.6.8) ^26^. Remaining reads were analyzed by applying the peak-calling algorithm MACS2 (v2.1.1) ^27^. Bigwig tracks were obtained for visualization on individual samples using deeptools (v3.1.0) ^28^. Differential ATAC-seq analysis was performed using DiffBind ^29^. Nearest genes were annotated using ChIPseeker 30. Transcription Factor motif enrichment analysis was performed using Bioconductor package LOLA (Locus overlap analysis or enrichment of genomic ranges; R package version 1.24.0) with RStudio (v3.6.1) ^31^.

#### Cell-line RNA-seq analysis

Cell line RNA-seq data was processed with three replicates. Paired-end reads were aligned to the human reference genome (hg19) using the STAR aligner with default parameters ^32^. Counts were obtained using featureCounts ^33^. Bigwig tracks were obtained for visualization on individual samples using deeptools (v3.1.0) ^28^. Downstream analysis including normalization and differential expression analysis was performed using DESeq2 ^34^. Genes were categorized as differentially expressed if abs(L2FC > 0.58, p-value < .05). Pathway analysis was performed using enrichR ^35^.

#### Primary patient sample RNA-seq analysis

Count data was generated using HTSeq ^36^. Downstream analysis including normalization and differential expression analysis was performed using DESeq2 ^34^. Genes were categorized as differentially expressed if abs(L2FC > 0.58, p-value < .05). Pathway analysis was performed using enrichR ^35^.

### ChIP-seq analysis

Cell line ChIP-seq data was processed with three replicates. Paired-end reads were aligned to the human reference genome(hg19) with Bowtie2 (v2.3.4.1) ^25^. Reads with low mapping quality <30 were discarded using Trimmomatic (v0.33; ref. 23) ^37^. Due to the broad/diffuse peaks created by H3K36me2 and H3K27me3, peaks for these marks were called by SICER that uses a cluster enrichment–based analysis ^38^. H3K27ac and H3K9ac peaks were called using MACS2 (–broad; ref. 30) ^27^. Bigwig tracks were obtained for visualization on individual samples using deeptools (v3.1.0) ^28^. Differential ChIP-seq analysis was performed using DiffBind ^29^. Peaks were categorized as differentially accessible if abs(L2FC > 1.0, FDR < .01). Nearest genes were annotated using ChIPseeker ^30^.

#### Hi-C analysis

Raw Hi-C sequencing data was processed with the hic-bench platform ^19^. Cell line Hi-C data was processed as single replicates. Data was aligned against the human reference genome(GRCh37/hg19) with bowtie2 (version 2.3.1) ^25^. The reads used for downstream analyses were classified as accepted intra-chromosomal read-pairs by the GenomicTools tools-hic filter command in the hic-bench platform. The number of accepted intra-chromosomal read-pairs varied between ~40 and ~140 million for all samples (Supplementary Fig. 1a). Interaction matrices for each chromosome separately were created by the hic-bench platform at 40kb resolution. Filtered read counts were normalized by iterative correction and eigenvector decomposition (ICE)^39^. To account for variances in read counts of more distant loci, distance normalization for each chromosome matrix was performed.

#### Compartment Analysis

The Cscore tool algorithm was used to assign active (A) and inactive (B) compartments ^20^. Differential Hi-C analysis with compartment scores was performed with two-sided t-tests.

#### Intra-TAD activity Analysis

Intra-TAD activity was assessed using the hic-ratio algorithm developed within hic-bench ^17,19^. Differential Hi-C analysis with TAD activity scores was performed with two-sided t-tests.

#### Integration of compartment data with other datasets

To show the correlation between the compartment changes and the various sequencing datasets, we mapped the peaks obtained from H3K36me2, H3K27me3, H3K27ac, ATAC-seq, and the genes obtained from RNA-seq to the AB, BA, and stable regions. We calculated the peak intensity fold change or gene expression fold change for peaks or genes assigned to a compartment region between NSD2 High and Low and showed the correlation with boxplots. Statistical significance was assessed using a paired two-sample t-test.

## Data Availability

All raw and processed data files generated through high-throughput sequencing for this publication (Hi-C) has been deposited in the National Center for Biotechnology Information Gene Expression Omnibus (GEO; http://www.ncbi.nlm.nih.gov/geo) and are accessible through GEO Series accession number (in progress). Previously generated high-throughput sequencing data (ChIP-seq, ATAC-seq, RNA-seq) used in this publication is accessible through GEO Series accession number GSE149159. Patient sample RNA-seq raw and processed data files acquired from COG Biobank have been deposited in the National Center for Biotechnology Information database of genotypes and phenotypes (dbGaP; https://www.ncbi.nlm.nih.gov/gap/) and is accessible through accession number (in progress).

## Acknowledgements

A.T. is supported by the NCI/NIH P01CA229086, NCI/NIH R01CA252239, NCI/NIH R01CA260028 and NIH/NCI R01CA140729. W.L.C has received grants from the National Cancer Institute of Health (R01CA140729 and R01CA260028), The Leukemia and Lymphoma Society Specialized Center for Research (7010-14), Perlmutter Cancer Center Arline and Norman M. Feinberg Pilot Grant for Lymphoid Malignancies, HHOW Scholar Hope Grant, and the Perlmutter Cancer Center (P30 CA016087). J.P. received funding from the St. Baldrick’s Foundation (Fellowship Award, 524986), Alex’s Lemonade Stand Foundation (Young Investigator Grant), and Pediatric Cancer Foundation (Fellowship Training Grant). We would like to thank the Genome Technology Center (GTC) for expert library preparation and sequencing, and the Applied Bioinformatics Laboratories (ABL) for providing bioinformatics support and helping with the analysis and interpretation of the data. GTC and ABL are shared resources partially supported by the Cancer Center Support Grant P30CA016087 at the Laura and Isaac Perlmutter Cancer Center. This work has used computing resources at the NYU School of Medicine High Performance Computing (HPC) Facility.

## Author Contributions

Conceptualization & study design, W.C. and A.T.; Investigation, S.N.; Formal analysis, S.N.; Writing—original draft, S.N.; Supervision, N.E., W.C. and A.T.; Funding acquisition, S.N., N.E., W.C., A.T.

## Competing Interests

The authors declare no competing interests.

## Notes

**CONFLICTS OF INTEREST**, The authors have no relevant affiliations or financial involvement with any organization or entity with a financial interest in or financial conflict with the subject matter or materials discussed in the manuscript.

### Competing Interest Statement

The authors have declared no competing interest.

