## Supplemental Files for "NSD2 E1099K drives relapse in pediatric acute lymphoblastic leukemia by disrupting 3D chromatin organization"

Supplementary Figure 1

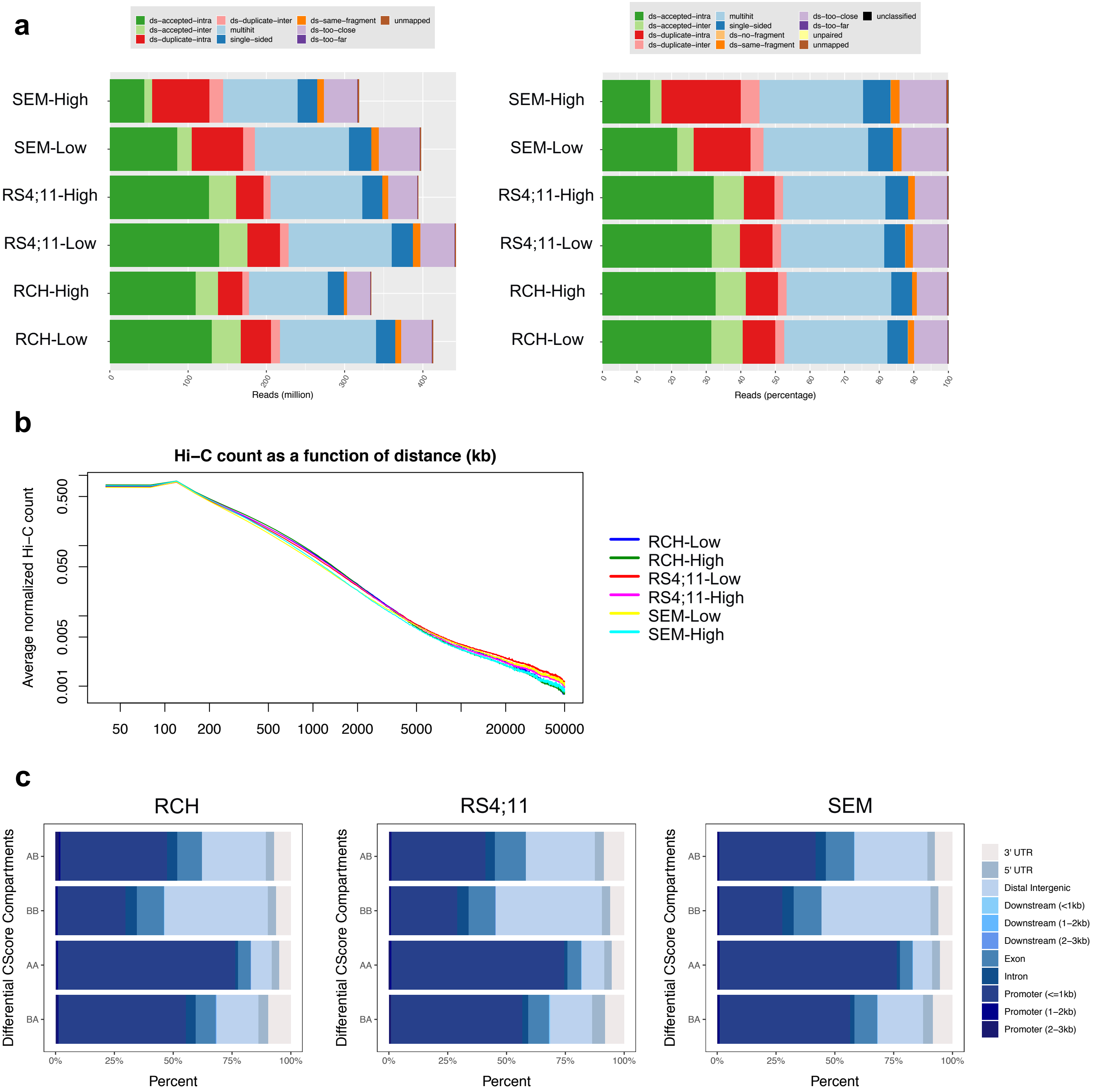

**Supplementary Figure 1. a.** Hi-C million read counts (left) and read percentage (right) per cell line. **b.** Average normalized Hi-C counts as a function of distance (kb) per cell line. c. Annotation of differential cscore compartments per cell line.

### Supplementary Figure 2

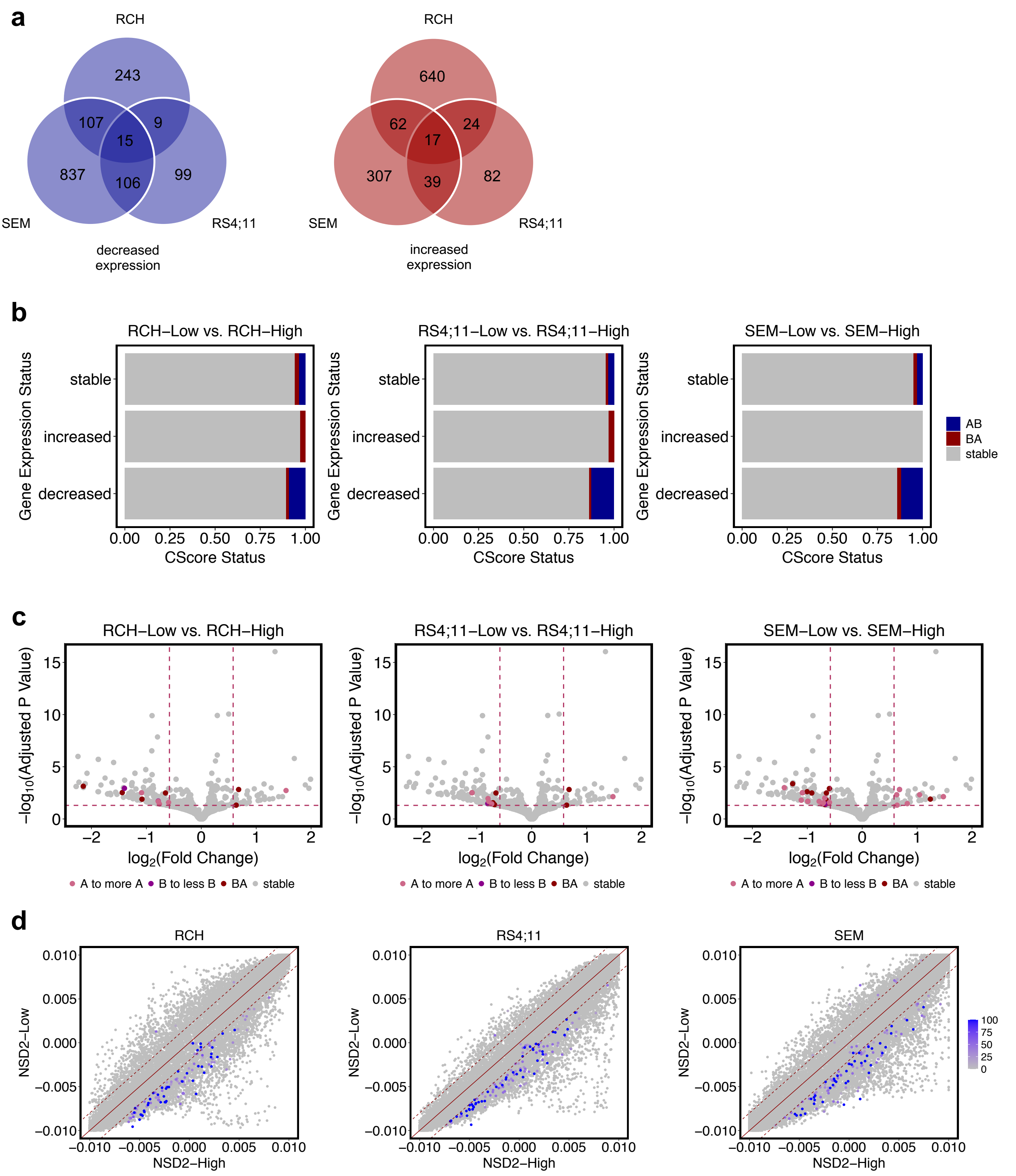

**Supplementary Figure 2. a**. Venn diagram showing overlap of differentially expressed genes (abs(L2FC) > 0.58, p-value < 0.05) between three B-ALL cell lines upon knockdown, decreased and increased genes (left and right respectively). **b.** Barplots showing fraction of compartment switches (AB, BA, stable) at differentially expressed genes(decreased, increased, stable). **c.** Volcano plots demonstrating differentially expressed genes (abs(L2FC) > 0.58, p-value < 0.05) highlighted by compartment switch or shift (A to more A, B to A, B to less B, or stable). **d.** Scatterplot demonstrating compartments colored by concordance score (percentage of genes with that compartment switch or shift that change in the same direction).

### Supplementary Figure 3

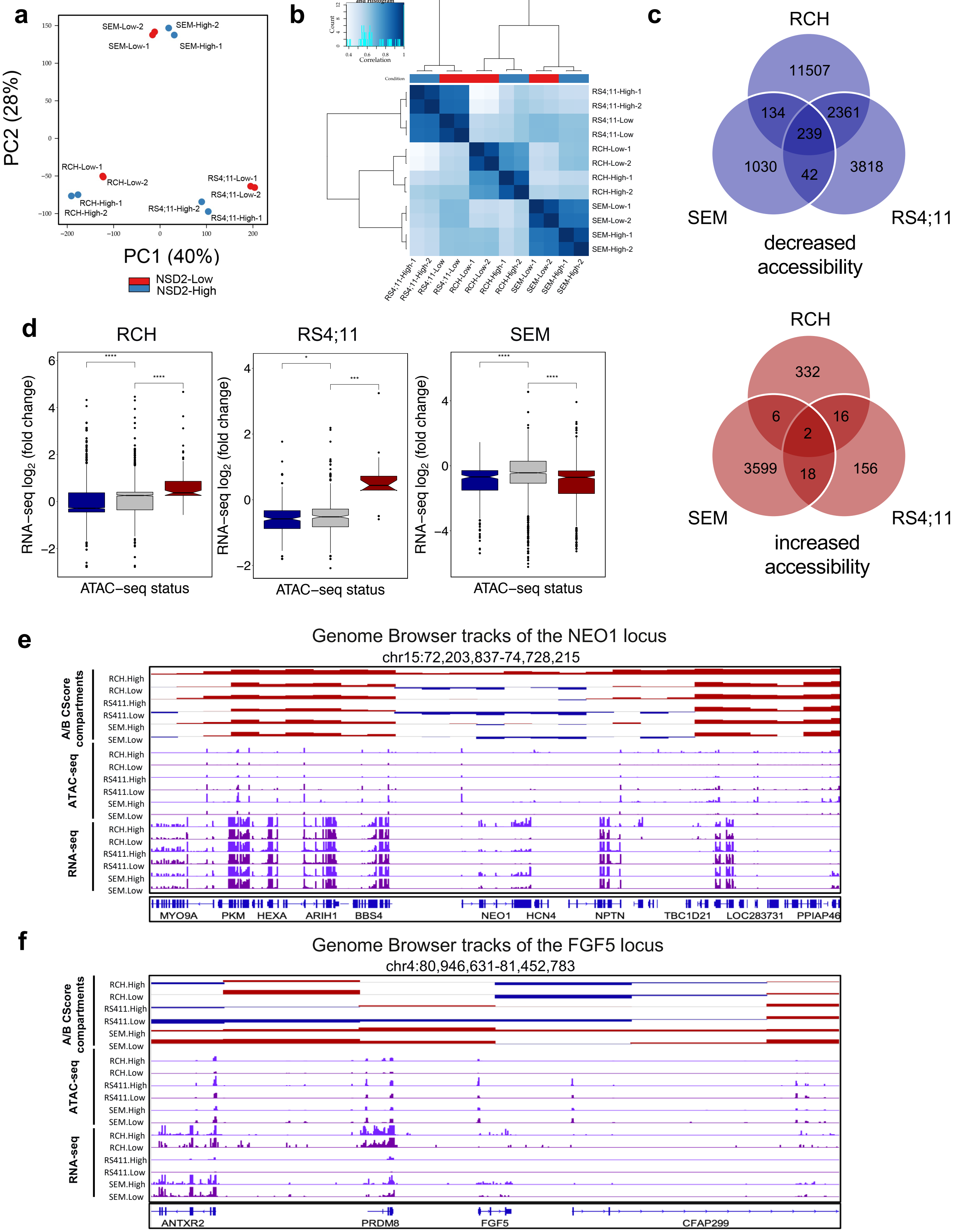

**Supplementary Figure 3. a.** PCA of ATAC-seq peaks for each cell line identifies three distinct clusters which are cell line-specific. **b.** Heat map representation of ATAC-seq results generated with DiffBind. **c.** Venn Diagram showing overlap of differentially accessible regions between three B-ALL cell lines downregulated and upregulated (top and bottom respectively; (FDR < 0.01, abs(log2(fold change)) > 1). **d.** Boxplots showing gene expression within ATAC-seq peaks that decrease, stable, and increase (blue, grey, and red) upon NSD2 knockdown for each cell line assessed with a two-sided t-test. **d.** Example of A to B compartment shift shared by all three cell lines at the NEO1 locus showing concordance with gene expression and chromatin accessibility. **e.** Example of a compartment shift specific to one cell line at the PRDM8 locus showing concordance with gene expression and chromatin accessibility.

### Supplementary Figure 4

**a**

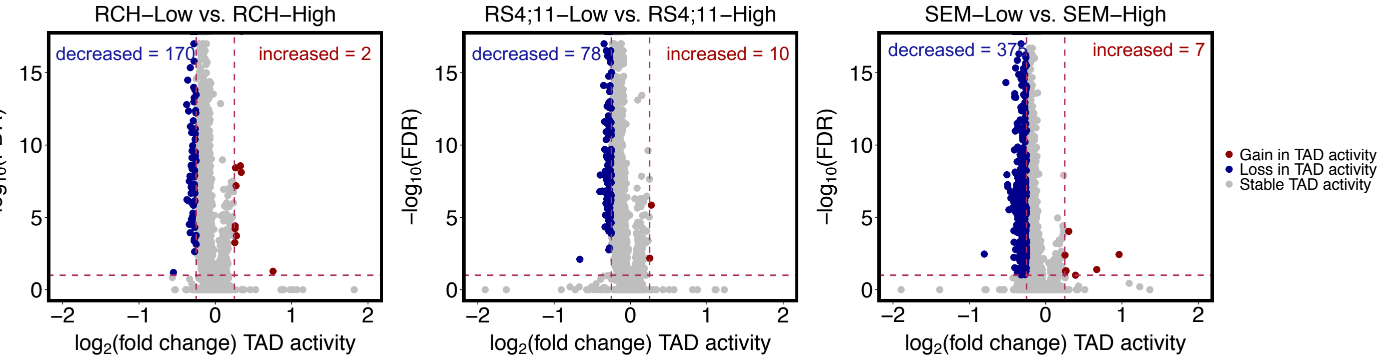


**b**

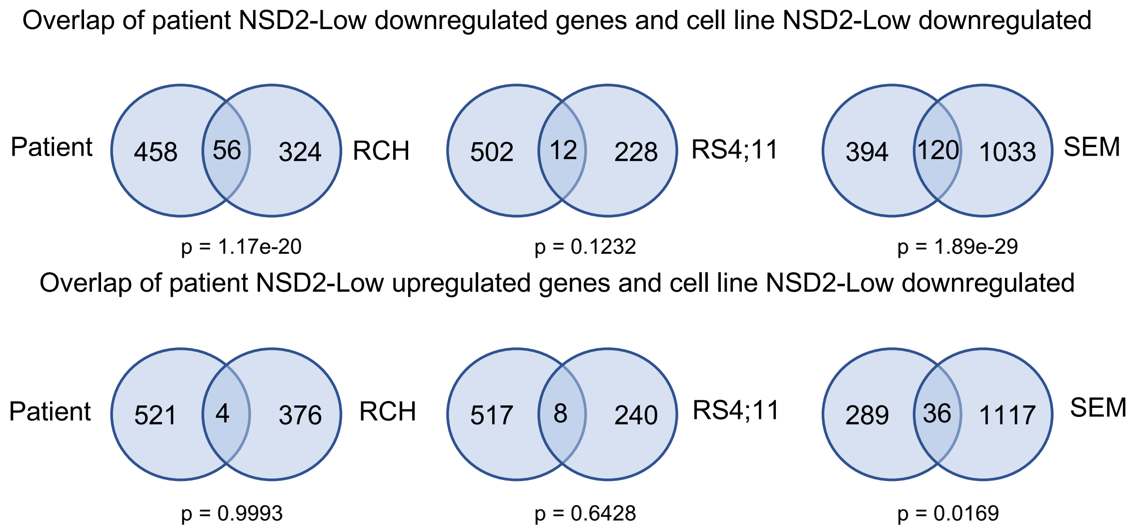


**c**

| Patient NSD2-Low down genes | Patient NSD2-Low down genes with AB Switch + shift | % |
|---|---|---|
| 514 | 195 | 37.94% |

**Supplementary Figure 4. a.** Volcano plots demonstrating differential intra-TAD activity upon knockdown per cell line (abs(L2FC) > 0.25, FDR < .01). **b.** Venn diagrams demonstrating significant overlap of patient NSD2-Low downregulated genes and cell line NSD2-Low downregulated genes and insignificant overlap of patient NSD2-Low upregulated genes and cell line NSD2-Low downregulated genes (one-tailed Fisher's exact test). **c.** Table presenting percentage of patient NSD2-Low downregulated genes overlapping a compartment switch/shift from any of the three cell lines.

### Supplementary Table 1

Explainability calculation for upregulated genes

| Cell line | #genes | Genes within BA Switch | explainability | Genes within Switch + Shift | explainability |
|---|---|---|---|---|---|
| RCH | 743 | 2 | 0.27% | 3 | 0.40% |
| RS411 | 162 | 2 | 1.23% | 3 | 1.23% |
| SEM | 425 | 0 | 0.00% | 8 | 1.88% |

Explainability calculation for downregulated genes

| Cell line | #genes | Genes within AB Switch | explainability | Genes within Switch + Shift | explainability |
|---|---|---|---|---|---|
| RCH | 374 | 20 | 5.34% | 69 | 18.45% |
| RS411 | 229 | 34 | 14.84% | 79 | 34.50% |
| SEM | 1065 | 27 | 2.54% | 74 | 6.95% |

**Supplementary Table 1.** Explainabillity calculations represent the amount of for upregulated and downregulated genes upon knockdown (abs(L2FC) > 0.58, p-value < 0.05) that can be explained by compartment switches and shifts.

### Supplementary Table 2

Patient Samples

| SJID | USI | Call_E1 | Call_S1 | chromosome | position | ref | mut | class | AAchange | fusion |
|---|---|---|---|---|---|---|---|---|---|---|
| SJALL045499 | PAVYIB | wt | som | chr4 | 1962801 | G | A | missense | E1099K | TCF3_PBX1 |
| SJALL057588 | PAWWLL | wt | som | chr4 | 1962801 | G | A | missense | E1099K | TCF3_PBX1 |
| SJALL068887 | PAXZXF | wt | som | chr4 | 1962801 | G | A | missense | E1099K | TCF3_PBX1 |

**Supplementary Table 2.** Table presents patient information associated with the three matched diagnosis-relapse pairs acquired from COG.

### Supplementary Table 3

Explainability calculation for ChIP-seq peaks

| PEAK | #peaks | AB Switch peaks | explainability | AB Switch + Shift peaks | explainability |
|---|---|---|---|---|---|
| H3K36me2 | 636 | 65 | 10.85% | 86 | 13.52% |
| H3K27me3 | 376 | 38 | 11.76 | 53 | 14.10% |
| H3K27ac | 178 | 16 | 9.62% | 27 | 15.17% |

**Supplementary Table 3.** Explainabillity calculations represent the amount of ChIP-seq peaks that can be explained by compartment shifts and switches.
